# Deep Learning Prediction of Glycopeptide Tandem Mass Spectra Powers Glycoproteomics

**DOI:** 10.1101/2024.02.03.575604

**Authors:** Yu Zong, Yuxin Wang, Xipeng Qiu, Xuanjing Huang, Liang Qiao

## Abstract

Protein glycosylation plays a significant role in numerous physiological and pathological cellular functions. Glycoproteomics based on liquid chromatography-tandem mass spectrometry (LC-MS/MS) studies the protein glycosylation on a proteome-wide scale to get combinational information on glycosylation site, glycosylation level and glycan structure. However, the current sequence searching-based methods for glycoproteomics often fall short in glycan structure determination due to the limited occurrence of structure-determining ions. While spectral searching methods can utilize fragment intensity information to facilitate the identification of glycopeptides, its application is hindered by the difficulties in spectral library construction. In this work, we present DeepGP, a hybrid deep learning framework based on Transformer and graph neural network (GNN), for the prediction of MS/MS spectra and retention time of glycopeptides. Two GNN modules are utilized to capture the branched glycan structure and predict glycan ions intensity, respectively. Additionally, a pre-training strategy is implemented to alleviate the insufficiency of glycoproteomics data. Testing on multiple biological datasets, we demonstrate that DeepGP can predict MS/MS spectra and retention time of glycopeptides closely aligning with the experimental results. Comprehensive benchmarking of DeepGP on synthetic and biological datasets validates its effectiveness in distinguishing similar glycoforms. Remarkably, DeepGP can differentiate isomeric glycopeptides using MS/MS spectra without diagnostic ions. Based on various decoy methods, we demonstrated that DeepGP in combination with database searching can significantly increase the detection sensitivity of glycopeptides. We outlook that DeepGP can inspire extensive future work in glycoproteomics.

## Introduction

Post-translational modifications (PTMs) tremendously increase the complexity of proteome^1^. Glycosylation, impacting over 50% of mammalian proteins, is one of the most significant PTMs, playing a crucial role in numerous physiological and pathological cellular functions^2^. Aberrations in protein glycosylation have been linked to a number of diseases, ranging from inflammatory conditions and diabetes to infectious diseases, neurodegenerative disorders such as Alzheimer’s and Parkinson’s diseases, as well as various forms of cancer^3^. A deeper exploration into the field of protein glycosylation can illuminate the intricate molecular mechanisms that precipitate these conditions^4^. The enhanced understanding could also pave the way for early diagnosis and innovative therapeutic interventions of the diseases^5, 6^.

In protein glycosylation, glycan molecules are usually attached to the side chains of asparagine (N-linked glycosylation) or serine/threonine (O-linked glycosylation) amino acid residues. Unlike biomacromolecules such as DNA, RNA and protein, the biosynthesis of glycans is non-template driven, leading to a plethora of structures that exhibit heterogeneity in linkage, number and composition of monosaccharides^7^. This heterogeneity results in the existence of multiple variants of glycopeptides, thereby posing substantial challenges in distinguishing the glycopeptide isomers^8, 9^. To date, liquid chromatography coupled tandem mass spectrometry (LC-MS/MS) is the leading technique in glycoproteomics research. In MS/MS, glycopeptides fragment ions, including b/y and c/z ions from the cleavage of peptide bonds and B/Y ions from the disruption of glycan bonds, can be generated by collision-induced dissociation (CID) and electron-transfer dissociation (ETD). Glycopeptides can be identified by combining the MS/MS with the molecular weight and the retention time information.

Various search engines, such as pGlyco series^10–12^, StrucGP^13^, MSFragger-Glyco^14^, O-Pair Search^15^, Byonic^16^, Glyco-Decipher^17^, GPSeeker^18^, etc., have been developed. These engines search the precursor and fragment mass-to-charge ratio (*m/z*) against the protein sequence and glycan structure database for intact glycopeptides identification. However, in many cases, *m/z* alone is insufficient for glycan structure determination due to the limited occurrence of structure-determining fragment ions^19^. As an alternative, spectra-match is also used for glycopeptide identification from LC-MS/MS data, wherein the experimental spectra are matched against a library of spectra to select the best matched candidates for glycopeptide identification, taking both intensity and *m/z* information into account. This approach overcomes the need for the observation of structure-determining fragment ions and improves identification sensitivity^20^. It is also computationally more efficient for high-throughput glycoproteomics studies^21^. GPQuest^20^ and GlycoSLASH^22^ have been developed for intact N-glycopeptide identification based on spectra-matching. Nevertheless, the construction of glycopeptide spectral libraries from synthetic or biological datasets presents a substantial challenge, due to the high expense associated with glycopeptide synthesis and the intricate complexity in characterizing glycopeptides from biological samples.

During the past years, deep learning has attracted significant attention in the prediction of peptide tandem mass spectra. Various deep learning-based spectra prediction tools, such as pDeep series^23–25^, Prosit^26^, DeepMass:Prism^27^, DeepDIA^28^, DeepPhospho^29^ and DeepFLR^30^, have been developed. Nevertheless, none of these tools can predict MS/MS spectra for glycopeptides. Taking pDeep2^23^ as an example, it poses good ability in the prediction of MS/MS spectra for peptides with various PTMs by encoding each PTM according to its atomic compositions. However, glycans with the same atomic compositions can have different structures and thus distinct chemical properties^6^. To predict the MS/MS spectra of glycopeptides, a novel deep-learning framework is required. This framework should be able to adeptly represent the branched architecture of glycans and to predict the intensity of the glycan fragment ions.

Here, we present DeepGP, a hybrid end-to-end deep learning-based framework for intact N-glycopeptides tandem mass spectra and retention time prediction. The deep learning framework consists of a pre-trained Transformer module and two graph neural network (GNN) modules. GNN is designed to directly capture graph structures, making it ideally suited for processing branched glycan structures. The two GNN modules of DeepGP are used to capture the glycan structures and to predict the glycan ions intensity, respectively. A pre-training strategy is exploited to alleviate the insufficiency of glycoproteomics data. We demonstrate the high accuracy of tandem mass spectra and retention time prediction by DeepGP using datasets of human and mouse samples. Furthermore, we demonstrate that the tandem mass spectra prediction by DeepGP can enhance the identification sensitivity of glycopeptides and even distinguish glycopeptide isomers for spectra without structure-determining ions.

## Results

### Construction and training of DeepGP

The model architecture of DeepGP is shown as **Fig. 1a**. DeepGP accepts glycopeptides as input and encodes multiple features of glycopeptides, including glycan structure, amino acid sequence, PTMs type (Glycosylation [N], Oxidation [M], Acetyl [Protein N-term] and Carbamidomethyl [C]), PTMs position and precursor charge state. The glycan structure is embedded by GNN, wherein the glycan is transformed into a graph with nodes representing monosaccharides and edges representing the linkages between the monosaccharides. The whole peptide is also treated as an anchor node (n_0_). This representation allows for encoding the glycan and generating an output matrix that captures both the glycan composition and the topological structure. In the embedding of peptide amino acid sequence, we add “[CLS]” as a special token before the sequence and “[SEP]” between any two amino acid tokens to simulate the fragmentation between amino acids. In a given analysis, the maximum padding length is determined by the longest peptide in a specific batch, and zero-padding is utilized to extend the length of sequences of the other peptides to match the maximal padding length. The up limit of padding length of 512-dimension is rooted in the configuration of the Bidirectional Encoder Representations from Transformers (BERT)^31^ utilized by our work. Other features of glycopeptides (PTMs type, PTMs position and precursor charge state) are also embedded into matrices of the same dimension as the amino acid sequence and the glycan. Subsequently, all the features are integrated through matrix addition for downstream prediction tasks.

**Fig. 1.**
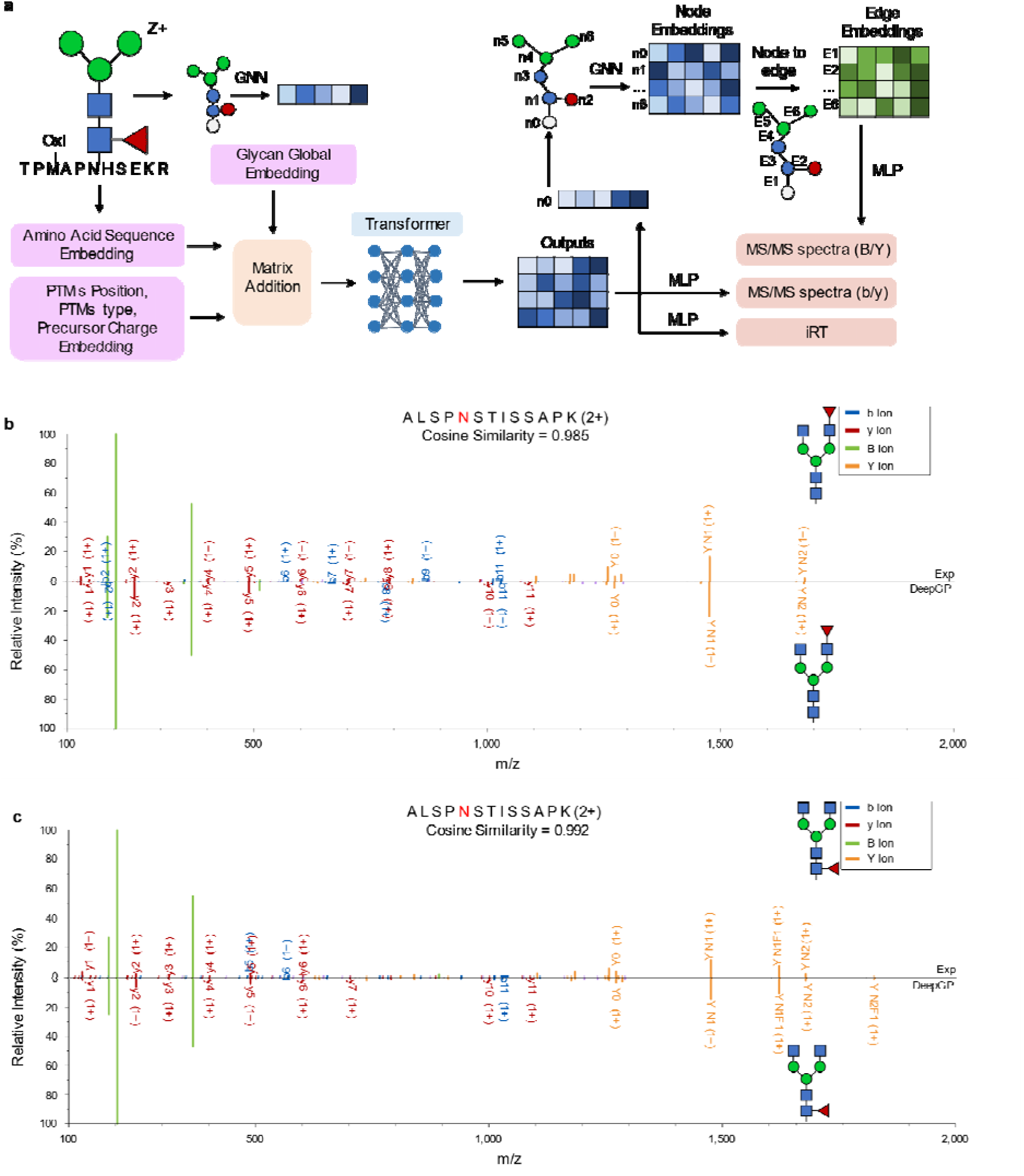
Model architecture and glycopeptide MS/MS spectra prediction. **(a)** The hybrid model architecture of DeepGP. Comparison of the experimental MS/MS spectra with the predicted MS/MS spectra of the glycopeptides with the same peptide sequence and glycan composition but with either **(b)** branched fucosylation or (**c)** core fucosylation. Top: the experimental MS/MS spectra; Bottom: the predicted spectrum. Assignments in common are labeled in color.

The learned glycopeptide representations are fed into a Transformer module for the prediction of MS/MS spectra and normalized retention time (iRT). An additional GNN module is used to predict the glycan fragment ions intensity. The Transformer outputs are pooled into a single vector that represents the embedding for the peptide node (n_0_). The embeddings of the remaining monosaccharide nodes are random-initialized. The GNN module is used to generate the node embeddings, which are then transformed into edge embeddings to predict glycan B/Y ions intensity through multilayer perceptron (MLP). The Transformer outputs are also used to predict b/y ions intensity and iRT through two other MLPs, respectively. In the MS/MS prediction, b/y and B/Y ions are considered concurrently in order to maintain the relative intensity of the b/y and B/Y ions. The iRT prediction is designed to be independent, enabling the use of optimal model parameters specific to iRT prediction. This also offers convenience for users who solely want to predict iRT or glycopeptide MS/MS spectra.

For MS/MS spectra, DeepGP predicts b/y fragment ions with the consideration of neutral loss of H_2_O and NH_3_. The b/y fragments with or without the monosaccharide HexNAc are both considered. Regarding B/Y ions, the neutral loss of H_2_O, NH_3_ and the monosaccharide Fuc are considered. Up to 16 types of B/Y fragment peaks or 24 types of b/y fragment peaks (**Supplementary Fig. 1**) are considered for each cleavage event by taking various neutral losses and charge states into consideration, as detailed in the **Methods** section. The total number of peaks is determined by the number of cleavage events and the number of ion types of each cleavage event. It is to be noted that ions deemed theoretically impossible aren’t manually excluded. Rather, predicting these ions incurs penalties in our scoring system, steering the model toward self-learning. Importantly, there is no restriction on the precursor charge states; while for fragment charges, we consider fragment charge up to +2 due to the rare occurrence of fragment ions with charge > 2.

We utilized stepped collision energy higher-energy collisional dissociation (SCE-HCD)^11,32^ MS/MS data for model training, validation and test, because the SCE-HCD method can generate numerous fragment ions for both glycan and peptide components. All the collected datasets were analyzed by pGlyco3^12^. Five SCE-HCD mouse datasets^11^ (Mouse_1-5, **Supplementary Table 1**) were used. These datasets encompass a total of 19,978 glycopeptide MS/MS spectra after the data processing as detailed in the **Methods** section, each associated with a distinct glycopeptide embedding pattern. The data were partitioned into training and validation data, employing a 9:1 ratio. DeepGP employs sqrt-cosine similarity (sqrt-COS, detailed in **Methods**) as the metric for MS/MS prediction, with the consideration that peak intensities follow a Poisson distribution and that the square root transformation can stabilize the peak intensity variance^21^. Besides, sqrt-COS can put more weight on low-intensity peaks and thus benefit the prediction of low-abundance ions.

We evaluate the performance of three GNN architectures, namely Graph Convolutional Network (GCN)^33^, Graph Isomorphism Network (GIN)^34^ and Graph Attention Network (GAT)^35^, for glycan embedding and B/Y ions intensity prediction. GCN utilizes convolutional operations on graphs to obtain node representations, and implements message-passing protocols to aggregate the representations of adjacent nodes^33^. GIN presents a simple yet expressive architecture, gaining notable distinction in the field of graph isomorphism testing^34^. GAT incorporates an attention mechanism, enabling the model to focus on the most relevant aspects of the inputs^36^. We tested the performance of the three GNN structures for either task of glycan embedding or glycan ions intensity prediction with the same hyperparameters and other model settings. Notably, in terms of glycan embedding, no performance differences were observed among the three GNN architectures (**Supplementary Fig. 2**). We chose GCN to represent the glycan structure for the following analysis in this work. For B/Y ions intensity prediction, GIN outperformed both GCN and GAT, and hence we selected GIN as the optimal GNN architecture for B/Y ions intensity prediction (**Supplementary Fig. 3**).

Due to the limited availability of glycopeptide MS/MS datasets, we utilized a pre-training strategy. We trained DeepGP based on Transformer without pre-training^37^, BERT^31^ and DeepFLR^30^. These models were of the same model configuration, but varied on the usage of pre-training. Transformer started with randomly initialized parameters, while BERT benefited from pre-training with unlabeled natural language. DeepFLR, a BERT-based framework, received additional training with phosphoproteome datasets. The result showed that the DeepFLR model can reach the highest median sqrt-COS while requiring the least epochs (**Supplementary Fig. 4**).

**Fig. 1b** and **Fig. 1c** show the prediction performance using two glycopeptides as examples. The two glycopeptides share the same peptide sequence (ALSPNSTISSAPK) and glycan composition (Hex(3)HexNAc(4)Fuc(1)), but different glycan structures, i.e. one with branched fucosylation (**Fig. 1b**) and another with core-fucosylation (**Fig. 1c**). The results demonstrate that the predicted spectra for both glycopeptides closely resemble their corresponding experimental spectra. Notably, the predicted spectra capture not only the m/z difference but also the intensity variation between the two isomeric glycopeptides for non-structure determining ions.

### Performance evaluation for glycopeptide MS/MS prediction

The performance of glycopeptide MS/MS prediction by DeepGP was firstly evaluated with the five datasets of five mouse tissues, including Brain, Kidney, Heart, Liver and Lung (Mouse_1-5, **Supplementary Table 1**), previously published by Liu et al.^11^. Each dataset served as the test dataset in turn, with the remaining four datasets providing the training data. DeepGP demonstrated remarkable accuracy in predicting glycopeptide MS/MS spectra, with the median cosine similarity (COS) between the predicted and experimental mass spectra surpassing 0.95 across all the datasets (**Fig. 2a** Intact). Upon narrowing our focus to the glycan B/Y ions, the accurate prediction of glycan B/Y ions intensity was also confirmed, dismissing the influence of peptide b/y ions in calculating the spectra similarity (**Supplementary Fig. 5**). Furthermore, we intentionally removed glycopeptides present in the training datasets from testing datasets. Despite this stringent exclusion, we observed only a marginal drop in the cosine similarity compared with the original results, amounting from 0 to 0.007 (**Fig. 2a** Test-only). We also assessed the experimental variability by calculating the COS of MS/MS spectra from the same glycopeptides repeated in the training and test datasets. The result demonstrated that DeepGP can predict glycopeptide MS/MS spectra with quality approaching the experimental replication (**Fig. 2a**). As for the different charge states of glycopeptides, DeepGP consistently maintained a high COS in MS/MS prediction (**Supplementary Fig. 6**). Beyond COS, we further explored the distribution of Pearson correlation coefficients (PCC) between the predicted and experimental spectra (**Supplementary Fig. 7**), and high median PCC was also demonstrated. We further trained DeepGP using all the five datasets, and tested the performance of MS/MS prediction on all the five datasets. Good performance of MS/MS prediction was also observed for whole spectra and the B/Y ions (**Supplementary Fig. 8**). The COS between predicted and experimental spectra of glycopeptides with different charge states (**Supplementary Fig. 9**) and the distribution of PCC between predicted and experimental spectra (**Supplementary Fig. 10**) were also presented when the training and testing were performed on all the five datasets, underscoring the model’s robust performance.

**Fig 2.**
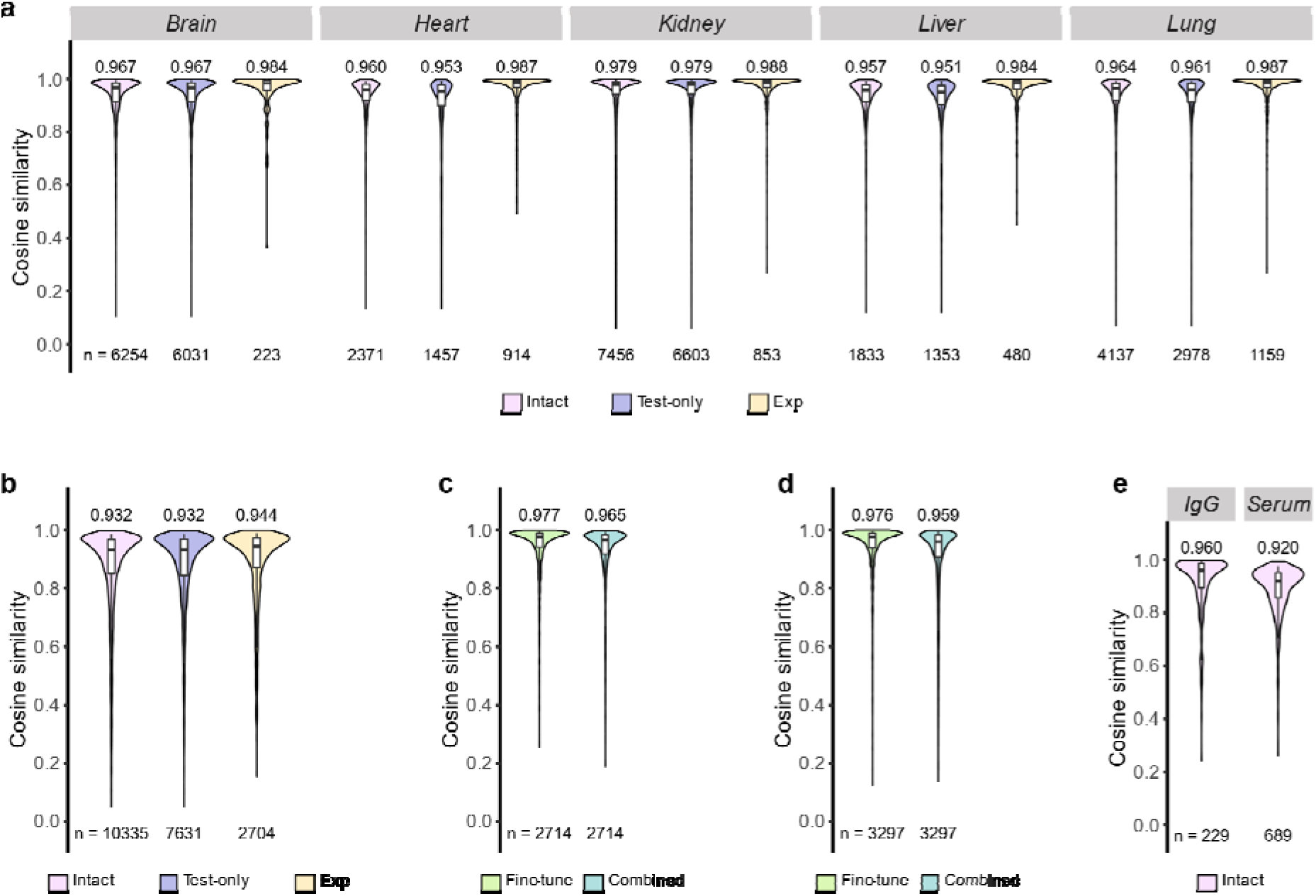
Performance of DeepGP in MS/MS spectra prediction. **(a)** The distribution of cosine similarity between the predicted and experimental spectra for the five mouse tissues datasets. Intact: cosine similarity computed for all glycopeptides within the test datasets. Test-only: cosine similarity computed for glycopeptides within the test datasets and not included in the training datasets. Exp: cosine similarity of repeatedly collected mass spectra in the training datasets and the test datasets of the same glycopeptides. **(b, c, d)** The distribution of cosine similarity between the predicted and experimental spectra on an external dataset Mouse_6. **(b)** DeepGP was trained using the five mouse tissue datasets (Mouse 1-5). **(c)** Mouse_6 was partitioned by “Run” into a training dataset (Run2, 3) and a test dataset (Run 1). DeepGP was fine-tuned using the training dataset based on the model trained with the five mouse tissue datasets (fine-tune) or trained by both Mouse_1-5 and the training dataset of Mouse_6 (combined). **(d)** Mouse_6 was divided into a training dataset (with HILIC enrichment) and a test dataset (with other enrichment). DeepGP was fine-tuned using the training dataset based on the model trained on the five mouse tissue datasets (fine-tune) or trained using both Mouse_1-5 and the training dataset of Mouse_6 (combined). **(e)** The distribution of cosine similarity between the predicted and experimental spectra on two Homo Sapiens datasets. The medians are indicated. The boxes and whiskers show the quantiles and 95% percentiles, respectively. The numbers of spectra for test are indicated below each graph.

We then applied the DeepGP trained by the five mouse tissue datasets to an external dataset^13^ (Mouse_6, **Supplementary Table 1**). Mouse_6 was generated by a different lab using a different mass spectrometer with different collision energies. Despite the different experimental conditions, DeepGP maintained a robust performance, yielding a median COS of 0.932, close to the COS (0.944 in median) between experimental MS/MS spectra from Mouse_6 and Mouse_1-5 (**Fig. 2b** Intact). Furthermore, we meticulously ensured that glycopeptides found in the training datasets (Mouse_1-5) were omitted from the testing dataset Mouse_6. Remarkably, we observed no change in cosine similarity, with a negligible difference of less than 0.001 (**Fig. 2b**, Test-only). To address the variance between datasets, fine-tuning was adopted to further enhance the performance of the MS/MS spectra prediction. The samples in the Mouse_6 dataset underwent three LC-MS/MS runs^13^, and we split the dataset by “Run” into the fine-tuning and test dataset. In detail, a subset of Mouse_6 (Run2, Run3) was leveraged for fine-tuning, while the remaining data (Run1) was allocated for testing (**Supplementary Table 1**). It turned out that fine-tuning led to a significant enhancement in prediction accuracy, with the median COS reaching 0.977 for the test dataset (**Fig. 2c**). We also trained DeepGP using the combination of the five mouse tissue datasets and a subset of Mouse_6 (Run2, Run3), and named the strategy as “combined learning”. It was observed that the fine-tuned model outperformed the combined learning model (**Fig. 2c**). The combined learning optimized the model to a validation dataset derived from all the six datasets, and therefore generating a model less optimized for Mouse_6 compared to the fine-tuned model. The Mouse_6 dataset also contained LC-MS/MS runs of glycopeptides enriched using hydrophilic interaction chromatography (HILIC) and mixed anion exchange (MAX) columns^13^. Therefore, we also partitioned the dataset as HILIC subset for fine-tuning and non-HILIC subset for test. In this scenario, the fine-tuning strategy also significantly enhanced the prediction accuracy, and outperformed the combined learning approach (**Fig. 2d**).

Finally, we evaluated DeepGP on human datasets. DeepGP was tested on an IgG dataset (Human_1)^38^ derived from Orbitrap Fusion Lumos with SCE-HCD at 20%-30%-40% and a serum dataset (Human_2)^39^ obtained from Q Exactive with SCE-HCD at 20%-25%-30%. Given that human datasets possess different glycans and peptides compared to the mouse datasets, we trained DeepGP with eight out of nine SCE-HCD human datasets (**Supplementary Table 1**) excluding the one for testing. DeepGP yielded a median COS of 0.960 for the IgG dataset and 0.920 for the serum dataset (**Fig. 2e**). These results reinforce the versatility of DeepGP across various organisms.

### Performance of DeepGP for iRT prediction

We also trained DeepGP for glycopeptide iRT prediction using the five mouse tissue datasets (Mouse_1-5). Each dataset was used as a test dataset in turn, with the remaining four datasets serving as the training data. Due to the variability of glycopeptide retention time values across datasets, the retention time values in the training datasets were calibrated to the test dataset using locally weighted scatterplot smoothing (LOWESS). The calibrated retention time values were then normalized to a range of 0-1 using the maximum retention time value within the datasets. PCC were calculated to assess the correlation between the predicted and experimental iRT values, resulting in coefficients ranging from 0.900 to 0.959 (**Fig 3a**). The interquartile ranges (IQRs) of the differences between the predicted and experimental iRT were below 0.07 (**Fig 3b**). These results demonstrate the good performance of DeepGP in glycopeptide iRT prediction. We also trained the DeepGP using all the five mouse tissue datasets, and tested on the mouse liver dataset (Mouse_4). In such a case the PCC can reach 0.978 (**Fig 3c**) with a IQR of 0.033 (**Fig 3d**).

**Fig 3.**
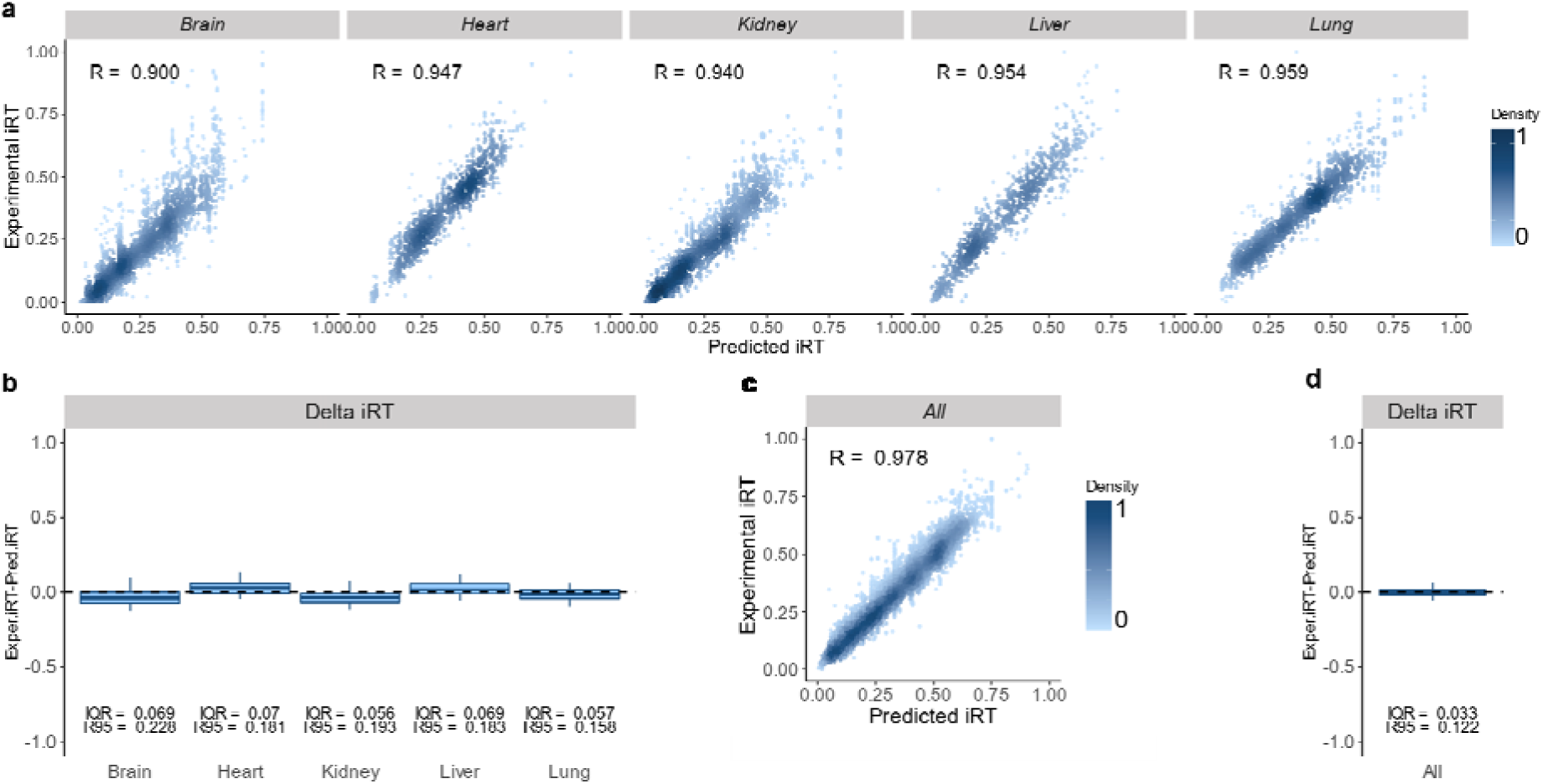
Performance of DeepGP in retention time prediction. **(a)** Correlation between the predicted and experimental iRT. Pearson correlation coefficients (R) is shown on the graph. Color gradation indicates relative density of data points. DeepGP was trained on four mouse datasets excluding the test dataset. **(b)** Differences computed between the predicted and experimental iRT. The boxes show interquartile ranges (IQR), and the whiskers show 95% percentiles; no outliers are shown. DeepGP was trained on four mouse datasets excluding the test dataset. **(c)** Correlation between the predicted and experimental iRT. DeepGP was trained on all the five mouse datasets, and tested on the mouse liver dataset. **(d)** Differences computed between the predicted and experimental iRT. DeepGP was trained on all the five mouse datasets, and tested on the mouse liver dataset.

### Differentiation of similar glycoforms by MS/MS prediction

Based on the accurate prediction of MS/MS for glycopeptides, we tried to perform spectra-matching-based glycopeptide identification. A synthetic dataset, including 14 glycopeptides with 7 peptide sequences and 2 sialylated glycans, was employed as the test dataset (**Supplementary Table 1**)^40^. The dataset was initially analyzed by pGlyco3. Subsequently, decoy glycopeptides were generated based on the identified glycopeptide, wherein the decoy glycopeptides had the same peptide sequences but substituted one NeuAc monosaccharide with two Fuc monosaccharides, resulting in a nominal mass difference of 1 Da. The glycan library consisted of nine glycans, including two target glycans and seven decoy glycans (**Supplementary Table 1**). DeepGP was trained by the six mouse (Mouse_1-6) and the nine human (Huam_1-9) datasets, and used to predict the MS/MS of both the identified glycopeptide and the corresponding decoy glycopeptides. The sqrt-COS was calculated between the experimental MS/MS spectra and the predicted MS/MS spectra of the corresponding target and decoy glycopeptides, and the match with the highest sqrt-COS was reported (**Fig 4**). Upon analyzing a total of 632 pGlyco3 identified MS/MS spectra of the synthetic dataset, 608 MS/MS spectra were still matched to the target glycopeptides, demonstrating the high resolving power of DeepGP in the differentiation of similar glycoforms. It should be noted that the spectra-matching method used in this analysis did not take into account the precursor mass difference.

**Fig 4.**
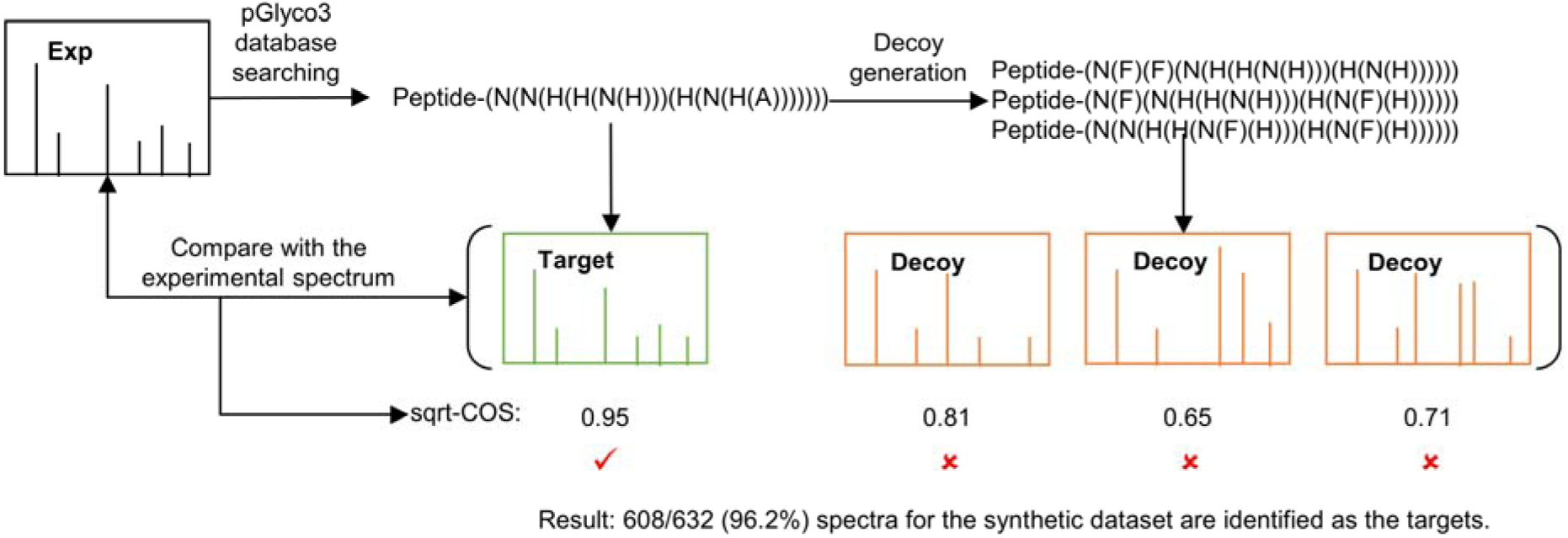
Deep-GP-based differentiation of similar glycoforms on a synthetic dataset. Decoy glycopeptides were generated by substituting one NeuAc monosaccharide with two Fuc monosaccharides, while retaining the same peptide sequences. DeepGP was employed to predict the MS/MS spectra for both the identified glycopeptide and the corresponding decoy glycopeptides. Re-identification results were reported based on the highest sqrt-COS between the predicted and experimental MS/MS spectra.

Furthermore, we benchmark DeepGP on two human datasets (Human_1, Human_2) using another decoy strategy. Given that NeuGc monosaccharides are typically absent in human samples, any human glycan that contains NeuAc and Hex monosaccharides can be assigned with a decoy glycan in the mouse library by substituting NeuAc and Hex with NeuGc and Fuc (**Supplementary Fig. 11**). The combined mass of NeuAc and Hex monosaccharides equals to that of NeuGc and Fuc. Both the human and mouse glycan libraries were from pGlyco3. It’s important to note that a single human glycan can correspond to multiple decoy glycans in the mouse glycan library containing NeuGc, wherein these decoy glycans share the same composition but different structures. The human datasets were firstly processed by pGlyco3, and the MS/MS spectra identified for glycopeptides containing NeuAc and Hex were collected. DeepGP was trained on eight of the nine human datasets, excluding the one for test, and used to predict the MS/MS spectra for both target and decoy glycopeptides. The sqrt-COS was calculated between the experimental MS/MS spectra and the predicted MS/MS spectra of the corresponding target and decoy glycopeptides, and the match with the highest sqrt-COS was reported. For the Human_1 dataset, 45/46 (97.8%) MS/MS spectra were still matched to the target glycopeptides. For the Human_2 dataset, 1018/1083 (94.0%) MS/MS spectra were still matched to the target glycopeptides. With all the benchmarking tests, it is demonstrated that the MS/MS prediction by DeepGP can be used to differentiate glycans with similar compositions.

### Identification of glycopeptides based on spectra-matching

With the ability to differentiate similar glycoforms, DeepGP was then applied to re-analyze the results by pGlyco3. Two mouse datasets of mouse brain (Mouse_1) and mouse liver (Mouse_4) were employed for the benchmarking. Alongside the final identification results, pGlyco3 also outputs a list of potential candidates for each glycopeptide MS/MS spectrum (pGlycoDB-GP-Raw*.txt). DeepGP was trained on four of the five mouse datasets, excluding the one for test, and used to predict MS/MS spectra for each candidate glycopeptide. The sqrt-COS was calculated between the experimental MS/MS spectra and the predicted MS/MS spectra, and the match with the highest sqrt-COS was reported. The identification results by DeepGP were then compared to those by pGlyco3. For the mouse liver dataset, out of all the DeepGP-reported results, 18,397/21,719 (84.7%) corresponded to the glycopeptides with the maximum “TotalScore” of pGlyco3 among all the candidate glycopeptides, when FDR was set as 1% during pGlyco3 analysis of the dataset (**Figure 5a**). Notably, the candidates with the highest “TotalScore” are not necessarily reported as the final identification results by pGlyco3, because multiple candidates can have the same “TotalScore”. When FDR was set as 100% during pGlyco3 analysis of the dataset, the proportion of DeepGP-reported results with the maximum “TotalScore” dropped to 69.1% (44,962 of 65,114) (**Fig. 5b**). Spectra identified at a higher FDR are generally less reliable than those at lower FDRs, and hence there was the drop during the DeepGP-based re-analysis. A similar analysis was conducted with the mouse brain dataset. In this case, 52.1% of all DeepGP-reported results (14,531 out of 27,868) had the maximum “TotalScore” at 1% FDR by pGlyco3 (**Fig. 5c**), and 47.2% (25,308 out of 53,580) at 100% FDR by pGlyco3 (**Fig. 5d**). For both the mouse brain and mouse liver datasets, there was a positive correlation between the sqrt-COS metric score of DeepGP and the score of pGlyco3 (**Supplementary Fig. 12**).

**Fig 5.**
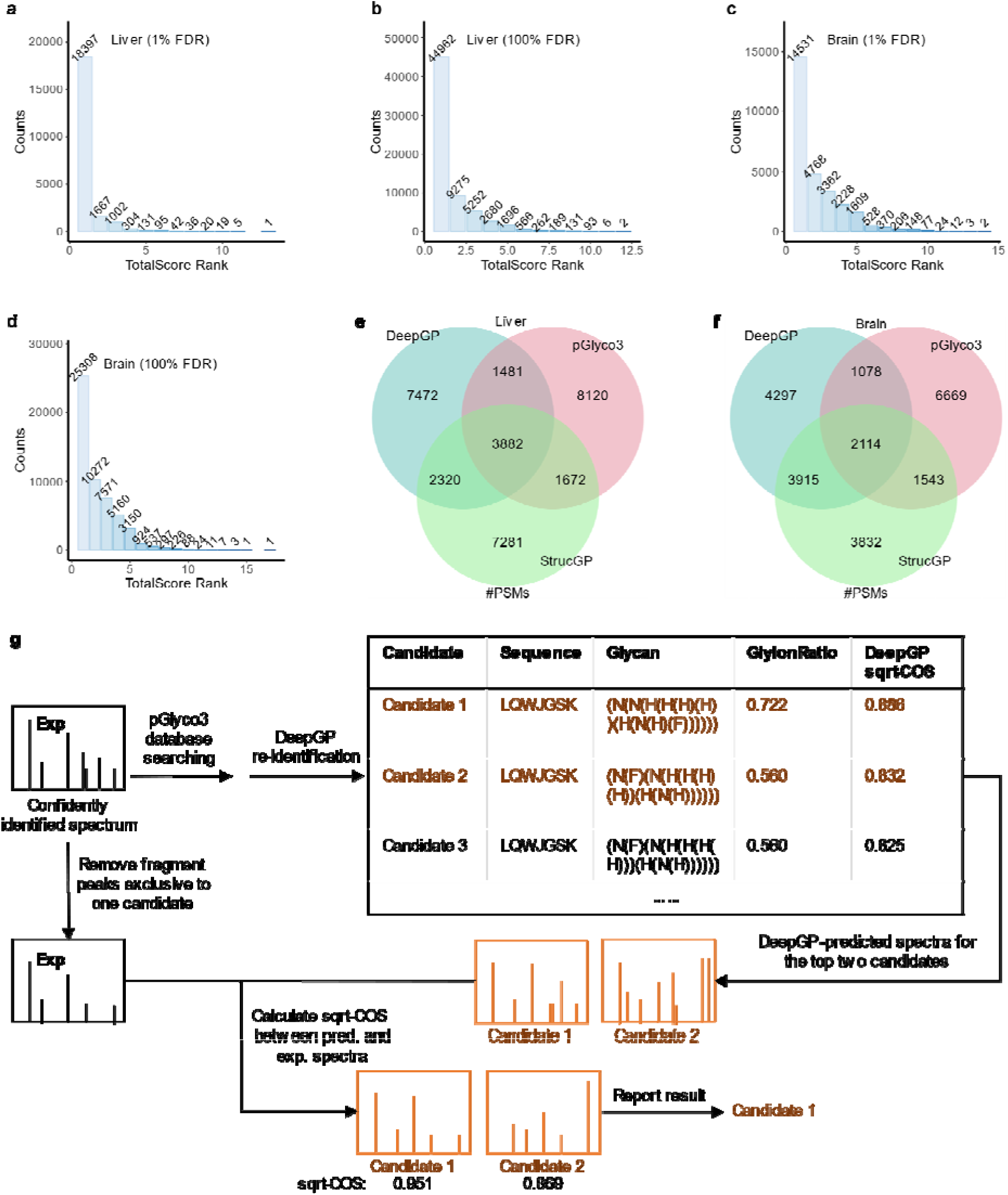
Performance of DeepGP in glycopeptide identification. Distribution of “TotalScore” ranks by pGlyco3 across all the identification results by DeepGP for the MS/MS spectra extracted from **(a)** the mouse liver dataset at 1% FDR of pGlyco3, **(b)** the mouse liver dataset at 100% FDR of pGlyco3, **(c)** the mouse brain dataset at 1% FDR of pGlyco3 and **(d)** the mouse brain dataset at 100% FDR of pGlyco3. Venn diagram of the number of PSMs identified by DeepGP, pGlyco3 and StrucGP based on **(e)** the mouse liver data and **(f)** mouse brain data. **(g)** Glycopeptide identification for spectra removing diagnostic ions.

Then, a comparative analysis of the identification results for the two mouse datasets (brain and liver) was performed utilizing DeepGP, pGlyco3 and StrucGP. The identification results from StrucGP were retrieved directly from the publication^13^. Only MS/MS spectra identified by both pGlyco3 and StrucGP with the same peptide amino acid sequence were extracted for comparison. Besides, the glycan identified by StrucGP should be present in pGlyco3’s glycan library, and the MS/MS spectra identified as multiple co-eluted glycopeptides were filtered. DeepGP was employed to search against glycopeptides identified by StrucGP and all the candidates by pGlyco3 using the same strategy aforementioned. As shown in **Fig. 5e**, **Fig. 5f**, there was a high discrepancy among the three methods in glycan structure identification, which is to date still an issue to be solved in glycoproteomics.

Glycopeptide identification through sequence searching inherently struggles to determine glycan structure when structure-determining ions are absent from the experimental spectra. However, multiple isomers may be distinguished via the spectra-matching. In this study, we tested DeepGP for the identification of glycan structure in the absence of diagnostic ions for the mouse liver dataset. MS/MS spectra with high-confidence identification results by pGlyco3 were collected. For the mouse liver dataset, there were 867 MS/MS spectra matching the criteria. For these spectra, the top two candidates with the highest sqrt-COS from DeepGP were retained. The diagnostic ions that can differentiate the two candidates were then removed from the experimental MS/MS, and the sqrt-COS was calculated again between the experimental MS/MS after removing diagnostic ions and the predicted MS/MS to report the result with a higher sqrt-COS (**Fig. 5g**). 842/867 (97.1%) MS/MS spectra maintained their original identification. The results demonstrate the robustness of DeepGP in the analysis of MS/MS spectra in the absence of diagnostic ions.

### DeepGP enhances the identification sensitivity of glycopeptides

We further performed rescoring by DeepGP-based spectra-matching to explore the value of DeepGP in glycoproteomics. A database entrapment method was adopted to evaluate the glycopeptide identification sensitivity by rescoring, referred to the work by Sun et al.^41^, Zeng et al.^12^ and Liu et al.^11^. Fission yeast glycoproteome datasets (Yeast_1, Yeast_2, **Supplementary Table 1**) were firstly searched by pGlyco3 against a large glycan database, which encompasses 4299 glycans, derived from the combined built-in resources of pGlyco-N-HighMannose.gdb and pGlyco-N-Human-multi.gdb of pGlyco3. The 4299 glycans contain a large portion of entrapment, as fission yeast should only contain high-mannose type glycans. All the glycoPSMs were reported without limiting FDR and ranked by the pGlyco3 TotalScore. Then, DeepGP was used to predict the MS/MS spectra of the identified glycopeptides, and the sqrt-COS was calculated between the predicted MS/MS spectra and the corresponding experimental MS/MS spectra. A combined score was then formulated by adding the normalized score (sqrt-cosine similarity) from DeepGP and the normalized score (TotalScore) from pGlyco3:

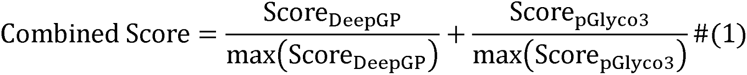

Based on the combined score, we re-ranked the identified glycoPSMs. For both rankings by pGlyco3 TotalScore and by the combined score, the glycan error rate was assessed by the ratio of identified glycoPSMs featuring glycans containing NeuAc (A) or Fucose (F), which are typically not anticipated in fission yeast samples. It should be noted that the ratio of entrapment glycoPSMs (decoy ratio) cannot be equal to the error rate or FDR, but a higher decoy ratio demonstrates a higher error rate.

Two fission yeast datasets were employed for the benchmarking, the Yeast_1 (**Supplementary Table 1**) originally reported by Yang et al.^40^ and the Yeast_2 (**Supplementary Table 1**) originally reported by Liu et al.^11^. The DeepGP models were initially trained on human samples, as detailed in **Supplementary Table 1**, with the exception of Human_5, a blend of human and budding yeast, and then fine-tuned with Yeast_1 or Yeast_2. The result demonstrated a notable enhancement in sensitivity at specific decoy ratios when DeepGP supplemented the identification by pGlyco3 (**Fig. 6**). With Yeast_1, the combination of DeepGP and pGlyco3 (DeepGP+pGlyco3) led to the identification of an additional 10,556 glycoPSMs at a decoy ratio of 5% (**Fig. 6a**). The Venn diagrams elucidated that DeepGP+pGlyco3 encompassed nearly all the PSMs (8614/8828=97.6%) (**Supplementary Fig. 13**) and glycopeptides (4766/4859 = 98.1%) (**Fig. 6b**) identified by pGlyco3, while also recognizing a significantly additional number of both PSMs (10770/8828=122.0%) and glycopeptides (8113/4859 = 167.0%). At a more stringent 1% decoy ratio, DeepGP+pGlyco3 identified 1,380 PSMs, in contrast to only 228 PSMs identified by pGlyco3. DeepGP+pGlyco3 successfully reported all the PSMs (**Supplementary Fig. 14a**) and glycopeptides (**Supplementary Fig. 14b**) identified by pGlyco3, while recognizing a significantly additional number of both PSMs (1152/228 = 505.3%) and glycopeptides (737/180 = 409.4%). Similar performance was observed with the Yeast_2 dataset (**Fig. 6c,d, Supplementary Fig.15, 16**).

**Fig. 6.**
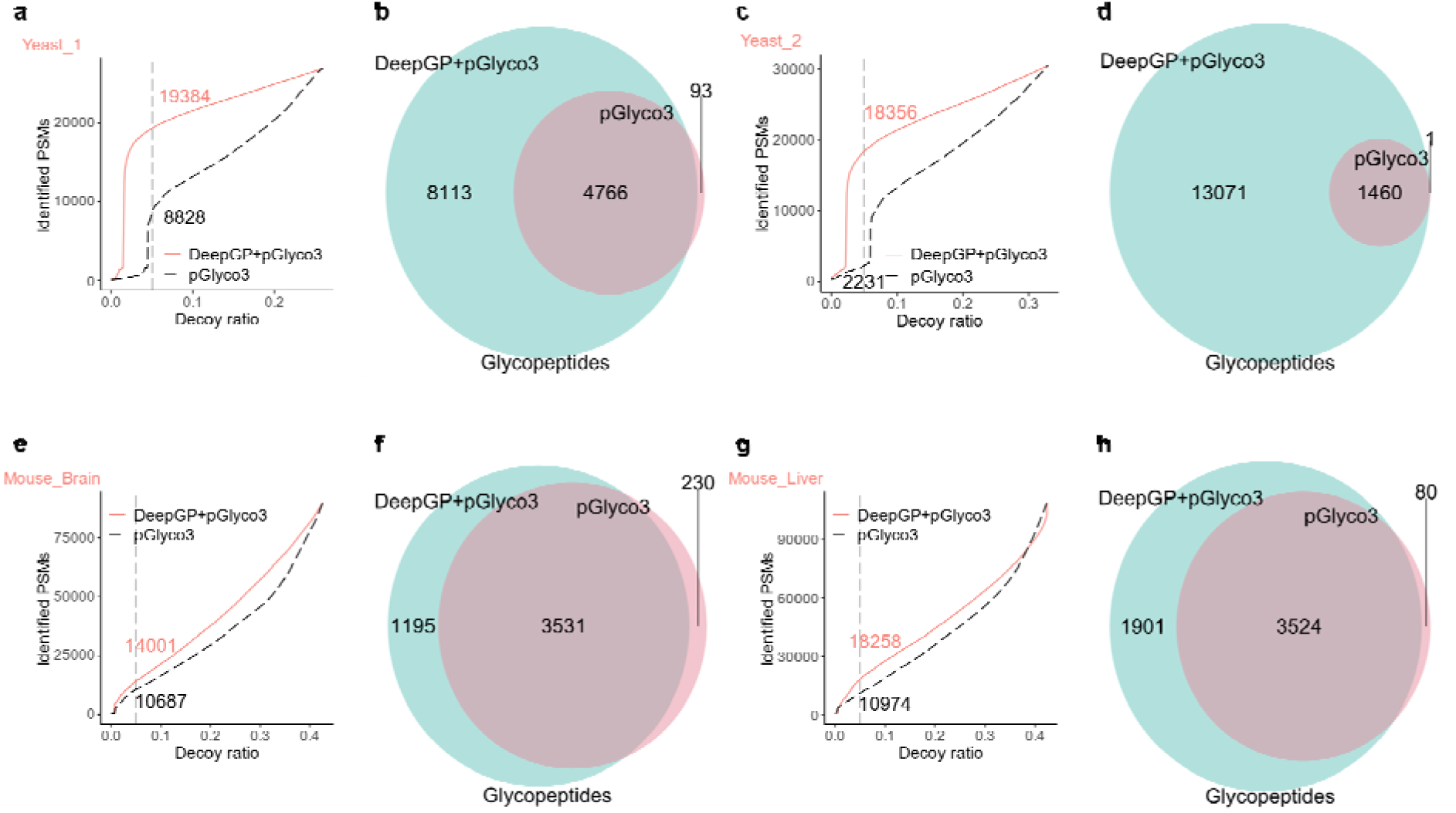
Comparison of the number of identified glycoPSMs under a given decoy ratio by DeepGP in combination with pGlyco3 (DeepGP+pGlyco3) and by pGlyco3 alone on **(a)** Yeast_1, **(c)** Yeast_2, **(e)** Mouse_Brain and **(g)** Mouse_Liver. The number of identified PSMs at 5% decoy ratio is indicated. Venn diagram of the number of glycopeptides identified by DeepGP+pGlyco3 and pGlyco3 for **(b)** Yeast_1, **(d)** Yeast_2, **(f)** Mouse_Brain and **(h)** Mouse_Liver at the decoy ratio of 5%.

Inspired by the work of Shen et al.^13^, and the thesis of Zhang^42^, we adopted another decoy method to evaluate the performance of DeepGP in the identification of glycopeptides. Different from the typical decoy methods generating decoy glycopeptides, the method generates decoy spectra by adding or subtracting random masses (1 to 30 m/z) to the peaks in the MS/MS spectra. To maintain the diagnostic ions for monosaccharides, for instance, Hex at the m/z of 366.139, we do not change the peaks with m/z < 367. Additionally, peaks with m/z exceeding the precursor are also retained. Furthermore, it is required to alter the m/z position of at least 20 peaks in an MS/MS spectrum. pGlyco3 was then applied to search the decoy spectra, as well as the corresponding original spectra. All the glycoPSMs were reported without limiting FDR and ranked by the pGlyco3 TotalScore. The combined scores were calculated in the same way aforementioned, and used to re-rank all the glycoPSMs. The error rate was assessed by the ratio of identified decoy spectra to the total number of spectra. It is also noted that the ratio of decoy spectra cannot be equal to the error rate or FDR, but a higher decoy ratio demonstrates a higher error rate. This ratio is also different from the entrapment ratio aforementioned.

The mouse brain (Mouse_1, **Supplementary Table 1**) and liver (Mouse_4, **Supplementary Table 1**) datasets were employed for the test. DeepGP models were trained on the other four mouse datasets when analyzing one mouse dataset. For the mouse brain dataset, DeepGP+pGlyco3 identified a larger number of glycoPSMs compared to pGlyco3 alone, as shown in **Fig. 6e**. At a 5% decoy spectra ratio, the combined approach identified 3314 more PSMs than pGlyco3. The Venn diagram (**Fig. 6f, Supplementary Fig. 17**) illustrates the PSMs and glycopeptides identified by DeepGP+pGlyco3 and pGlyco3 alone at the decoy spectra ratio of 5%. DeepGP+pGlyco3 identified 92.3% (9866/10687) of the PSMs identified by pGlyco3 alone, and identified 38.7% (4135/10687) additional PSMs. At the glycopeptide level, DeepGP+pGlyco3 identified 93.9% (3531/3761) of the glycopeptides identified by pGlyco3 alone, and identified 31.8% (1195/3761) additional glycopeptides. A similar performance was observed with the mouse liver dataset (**Fig. 6g,h** and **Supplementary Fig. 18**).

## Discussion

Glycoproteomics is a critical yet burgeoning field. The number of currently available glycoproteomics datasets is much fewer than datasets for proteins without focus on PTMs or with focus on other PTMs such as phosphorylation. Additionally, the lack of a standardized protocol for generating glycopeptide MS/MS data further diminishes the availability of suitable data for deep-learning model training. Diverse dissociation methods, such as ETD, CID, HCD, SCE-HCD, and electron-transfer/higher-energy collisional dissociation (EThcD), are employed in glycoproteomics. Even within a single method, like SCE-HCD, the applied stepped collision energies can vary. The insufficiency in datasets and diversity in protocols limit the width of knowledge that a deep learning model can acquire.

In addition to the limitations imposed by the availability of data, a community study indicates that different searching software or even varying searching settings can yield dramatically divergent identification results for glycoproteomics^43^. Current search software struggles with accurately identifying the glycan structure of glycopeptides. Although deep learning can tolerate a certain degree of inaccuracy, the challenges associated with glycopeptide identification could introduce noise into the training data and undermine the performance of DeepGP.

In order to alleviate the above-mentioned difficulties, DeepGP employs a pre-training strategy to mitigate data scarcity and utilizes GNN to process glycan structure. Pre-training involves training a deep learning model on a separate task before fine-tuning it to the target dataset^44^. We have demonstrated that pre-training can combat the shortage of data and enhance the performance of DeepGP. GNN is used to represent the branched structure of glycans and predict the intensity of glycan B/Y ions. In comparison to common deep neural network, GNN embedding is dependent on the graph structure, and hence can differentiate glycans with the same compositions but different linkages among monosaccharides. With 6 mouse glycoproteomics datasets and 9 human glycoproteomics datasets based on the SCE-HCD method, we demonstrated the good performance of DeepGP in MS/MS and iRT prediction. We only trained DeepGP to represent N-glycopeptides due to the even smaller amount of data available for O-glycopeptides. Given sufficient training data for O-glycoproteomics in the future, we foresee the potential of DeepGP to be applied to O-glycopeptides. We also anticipate that this approach can improve deep learning-based models for other biological molecules, such as cross-linked peptides and glycolipids, due to the power of GNN in representing structures.

Benchmarked on both synthetic and biological datasets, DeepGP has been demonstrated with the ability to differentiate similar glycoforms. Additionally, we present the identification results by DeepGP-based spectra-matching and compare the results of pGlyco3, StrucGP and DeepGP. Notably, DeepGP is able to distinguish isomeric glycopeptides without diagnostic ions. DeepGP can also elevate the identification sensitivity of glycopeptides by rescoring the results from database searching tools, such as pGlyco3, demonstrated with different decoy methods^11–13, 41, 42^. It should be emphasized that despite the decoy methods are widely adopted by many works the decoy ratio cannot be viewed as the real error rate or FDR. Due to the lack of large-scale synthetic glycopeptide datasets, it is still hard to determine the optimized decoy generation method and calculate the empirical FDR at the glycan level or the glycopeptide level. In this work, we use the decoy methods to compare under the same condition the performance of glycopeptide identification with or without DeepGP-based MS/MS prediction. DeepGP can particularly be valuable in producing MS/MS spectra that are challenging or even impossible to be obtained by experiments, such as decoy glycopeptide MS/MS spectra. This capability could facilitate the estimation of FDR of glycan structure in the future.

To date, there are a few works available on the prediction of glycopeptide MS/MS. Zhang reported glycopeptide MS/MS prediction by deep learning in the Master degree thesis^42^. Klein et al.^45^ and Zhang et al.^46^ reported glycopeptide MS/MS prediction based on kinetic models. As there is a lack of available code or sufficient guidance to replicate these works, we can only directly compare the reported results. As stated in Zhang’s thesis^42^, the deep learning model was trained on the four mouse tissues datasets other than the mouse brain dataset, and tested on the mouse brain dataset. A median cosine similarity of 0.943 was obtained. In our work, we adopted the same datasets for training and test (**Fig. 2a** Brain), and obtained a median cosine similarity of 0.967. The performance of the models by Klein et al.^45^ and Zhang et al.^46^ is worse than that of Zhang’s thesis^42^, and hence also worse than our work. It is important to note that the model reported by Zhang’s thesis^42^ predicts only glycan fragments and only ions with a single charge. In contrast, DeepGP predicts fragment charges of 1 and 2 for both peptide and glycan fragmentation, and considers a number of neutral loss and other modifications, such as Oxidation (M).

In summary, DeepGP, a hybrid end-to-end deep learning framework based on GNN and Transformer, has been developed for the prediction of glycopeptide MS/MS and retention time. The model architecture of DeepGP empowers it to capture glycan linkage heterogeneity along with other glycopeptide features including amino acid sequence, PTMs type, PTMs position and precursor charge state. DeepGP has shown potential in distinguishing similar glycoforms and even isomeric glycopeptides without diagnostic ions. Besides, DeepGP can enhance the identification sensitivity of glycopeptides by rescoring the identification results from database searching. With the development of glycoproteomics, particularly in mass spectrometry methods and data analysis software solutions, we anticipate that deep learning frameworks like DeepGP can find more applications in the future. We anticipate that DeepGP will enhance our understanding of the heterogeneity and complexity of glycoproteomics, and further aid biologists in the study of glycobiology.

## Methods

### Glycoproteome data analysis by pGlyco3

Data analysis was conducted using pGlyco3 (release: pGlyco3.0_build20210615, https://github.com/pFindStudio/pGlyco3/releases). The parameters for pGlyco3 included Oxidation (M) and Acetyl (Protein N-term) as variable modifications and with Carbamidomethyl (C) as a fixed modification. For the human and mouse datasets, we used the respective glycan databases embedded within pGlyco3, pGlyco-N-Human.gdb and pGlyco-N-Mouse.gdb respectively. For the fission yeast datasets, we used the combined glycan databases of pGlyco-N-HighMannose.gdb and pGlyco-N-Human-multi.gdb. The synthetic glycopeptide dataset was searched using the glycan database provided by the corresponding publication^40^. The precursor tolerance was set at 5 ppm, fragment tolerance at 20 ppm, and glycopeptide FDR at 0.01. Other parameters were left at default settings. FASTA files were from UniProt H. sapiens reference proteome (20,600 entries), UniProt M. musculus reference proteome (17,082 entries) and UniProt S. pombe reference proteome (5140 entries). For the synthetic glycopeptide dataset (Syn_1), FASTA file was compiled from synthetic glycopeptide sequences as stated in the original paper^40^.

### DeepGP model architecture

DeepGP is a hybrid deep-learning model consisting of a Transformer module and two GNN modules. DeepGP accepts glycopeptides as input and encodes multiple features of glycopeptides, including glycan structure, amino acid sequence, PTMs type (Glycosylation [N], Oxidation [M], Acetyl [Protein N-term], Carbamidomethyl [C]), PTMs position and precursor charge state. To embed the global structure of glycans, a 7-layer GCN is used. After embedding, the learned glycopeptide representations are fed into the Transformer for the prediction of MS/MS spectra or iRT. The pre-trained Transformer module parameters are sourced from DeepFLR, a BERT model trained on fourteen HCD phosphoproteome datasets, comprising over 467,000 PSMs^30^. The pre-trained Transformer module consists of 12 layers, each with two sublayers. The first sublayer is a multiheaded self-attention layer, and the second is a fully connected point-by-point feed-forward network. Residual connection and normalization are used in each sublayer. After the Transformer encoder, a 7-layer GIN is used to predict the glycan B/Y ions intensity. MLPs are used to transform the hidden states into predicted MS/MS. The model architecture can be easily modified using keywords. For instance, inputting “GNN_global_ablation=GIN” changes the GNN architecture for glycan global representation into GIN. More details for the usage of DeepGP are available in the “User Guide” file along with the code.

The model is optimized using Mean Square Error (MSE) loss. The MS/MS spectra prediction performance is assessed by sqrt-cosine similarity (sqrt-COS):

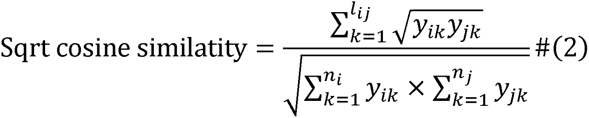

where *l_ij_* is the number of common peaks in spectrum *i* to spectrum *j*, *n_i_* and *n_j_* are the number of peaks recognized as glycopeptide fragments in spectrum *i* or spectrum *j*, respectively, and *y* is the intensity of the peak. A tolerance of 20 ppm was set to recognize the common peaks. If multiple peaks appear within the tolerance, the peak with the shortest distance was chosen. iRT prediction is assessed by coefficient of determination (R2) as implemented in the sklearn.metrics.r2_score (www.scipy.org). Hyperparameters are available in the code’s utilities (utils.py) of DeepGP.

DeepGP considers 24 types of b/y fragment peaks (b1, b1n, b1o, b1h, b1nh, b1oh, y1, y1n, y1o, y1h, y1nh, y1oh, b2, b2n, b2o, b2h, b2nh, b2oh, y2, y2n, y2o, y2h, y2nh, y2oh). The first character means ion type (b ions or y ions), followed by fragment charge (1 or 2). The final character(s) means neutral loss type (“o” means loss of H_2_O, “n” means loss of NH_3_, and “h” means loss of monosaccharide HexNAc). In our presentation, the peaks without a symbol “h”, (b1, b1n, b1o, y1, y1n, y1o, b2, b2n, b2o, y2, y2n, y2o) are b/y fragments with one HexNAc moiety, and the ones with a symbol “h” (b1h, b1nh, b1oh, y1h, y1nh, y1oh, b2h, b2nh, b2oh, y2h, y2nh, y2oh) are b/y fragments losing the HexNAc moiety. This nomenclature is to ensure that all the symbols unambiguously represent losses for convenience in coding. DeepGP considers 16 types of B/Y fragment peaks (B1, B1n, B1o, B1f, Y1, Y1n, Y1o, Y1f, B2, B2n, B2o, B2f, Y2, Y2n, Y2o, Y2f). The first character means ion type (B ions or Y ions). “f” means loss of monosaccharide Fuc, and the others follow the same nomenclature rules.

### DeepGP model training

DeepGP uses the search results by pGlyco3 for model training, validation and test. For compatibility purposes, DeepGP uses the nomenclature of monosaccharides defined by the pGlyco series. The nomenclature is listed in **Supplementary Table 2**. The training datasets were selected based on the organisms of the test datasets, and the test datasets were excluded from the training datasets unless explicitly stated. For model training, the combined datasets were split into training and validation datasets at a 9:1 ratio. Each spectrum in the training or test datasets displayed a unique embedding pattern (peptide sequence, glycan, PTMs type, PTMs position, and precursor charge). During model training, the MS/MS spectra intensity was normalized by the highest peak in the spectrum. The RT values of the glycopeptides in the test datasets served as retention time anchors for calibrating the retention time of the training datasets. We employed LOWESS to calibrate the RT values. Subsequently, these calibrated retention times were normalized to a range of 0-1 by the maximum retention time value within the dataset. Training of DeepGP on the data of five mouse tissues took 3 hours on a single RTX 3090 GPU with 150 epochs, a batch size of 256, and a learning rate of 1×10^−4^. It took less than 2 seconds to predict 40 spectra on a single RTX 3090 GPU.

### DeepGP prediction evaluation

The experimental spectra were converted to Mascot generic format (.mgf) using MsConvert from the ProteoWizard Package (3.0.11579). Predicted spectra were generated by DeepGP. A tolerance of 20 ppm was set to recognize the common peaks between the experimental and the predicted spectra. If multiple peaks appear within the tolerance, the peak with the shortest distance was chosen. Discrepancies between compared spectra resulted in the missing peak intensity being set as 0, followed by the calculation of the spectrum similarity. DeepGP offers several metrics, including square root cosine similarity (sqrt-COS), cosine similarity (COS), and Pearson correlation coefficient (PCC).

### Implementation and visualization

DeepGP was developed using python (3.8.3, Anaconda distribution version 5.3.1, https://www.anaconda.com/) with the following packages: FastNLP (0.6.0), pytorch (1.8.1), torchinfo (1.7.1), transformers (4.12.5), dgl (1.0.1), bidict (0.22.0), pandas (1.0.5) and numpy (1.18.5). Visualization was performed using custom scripts in R (4.0.2) with the following packages: VennDiagram (1.6.20) and ggplot2 (3.3.2). The mirrored spectra are plotted by GP-plotter (1.0.0, https://github.com/DICP-1809/GP-Plotter).

## Data availability

The raw datasets used in this study are available in the PRIDE^47^ database under accession code PXD005411^11^, PXD005412^11^, PXD005413^11^, PXD005553^11^, PXD005555^11^, PXD025859^13^, PXD015360^38^, PXD009654^39^, PXD023980^40^, PXD016428^48^, PXD005931^49^, PXD025455^50^, PXD009716^51^ and PXD005565^11^. Further details regarding the datasets and raw files used in this study can be found in **Supplementary Table 1**.

## Acknowledgements

We thank Prof. Mingliang Ye and Dr Zheng Fang from Dalian Institute of Chemical Physics, Chinese Academy of Sciences for assisting us in using GP-plotter.

## Author contributions

Y.Z. did the majority of coding work and data analysis, and wrote the original draft of the manuscript. Y.W. built the deep-learning model. X.Q. and X.H. assisted in the building of the deep-learning model. L.Q. finalized the manuscript and supervised all aspects of the work. All authors were involved in the design of this work.

