## Supplementary Information 1 for "Deep Learning Prediction of Glycopeptide Tandem Mass Spectra Powers Glycoproteomics"

Zong et al.

**Supplementary Table 1.** Datasets for deep learning model training, evaluation and test.

| Name | Sample | Instrument | Accession | Description | |
|---|---|---|---|---|---|
| Mouse_1 | Mouse (Brain) | Orbitrap Fusion Tribrid (Fragmentation: SCE-HCD 20%-30%-40%) | PXD005411[1] | 6254 unique glycopeptides* | Combined for training: 19,978 unique glycopeptides* |
| Mouse_2 | Mouse (Kidney) | | PXD005412[1] | 7456 unique glycopeptides* | |
| Mouse_3 | Mouse (Heart) | | PXD005413[1] | 2371 unique glycopeptides* | |
| Mouse_4 | Mouse (Liver) | | PXD005553[1] | 1833 unique glycopeptides* | |
| Mouse_5 | Mouse (Lung) | | PXD005555[1] | 4137 unique glycopeptides* | |
| Mouse_6 | Mouse | Orbitrap Fusion Lumos (0.7 m/z for 20% collision energy and 2 m/z for 33% collision energy) | PXD025859[2] | 10,335 unique glycopeptides* | Split by "Run":<br>Train (Run 2, 3): 7621 unique glycopeptides*<br>Test (Run 1): 2714 unique glycopeptides*<br>Split by "HILIC":<br>Train (enrich by HILIC): 7038 unique glycopeptides*<br>Test (enrich not by HILIC): 3297 unique glycopeptides* |
| Human_1 | Human (IgG) | Orbitrap Fusion Lumos (Fragmentation: SCE-HCD 20%-30%-40%) | PXD015360[3] | 229 unique glycopeptides* | |
| Human_2 | Human (Serum) | Q Exactive (Fragmentation: SCE-HCD 20%-25%-30%) | PXD009654[4] | 689 unique glycopeptides* | |
| Human_3 | Human (Serum & Urine) | Orbitrap Fusion Lumos (Fragmentation: SCE-HCD 20%-30%-40%) | PXD015360[3] | 17,104 unique glycopeptides* | |
| Human_4 | Human | Orbitrap Fusion Tribrid (Fragmentation: SCE-HCD 20%-30%-40%) | PXD023980[5] | 4251 unique glycopeptides* | |
| Human_5 | Human & Budding Yeast | Orbitrap Fusion Tribrid (Fragmentation: SCE-HCD 20%-30%-40%) | PXD023980[5] | 1953 unique glycopeptides* | |
| Human_6 | Human | Orbitrap Fusion Lumos (Fragmentation: SCE-HCD 20%- | PXD016428[6] | 4914 unique glycopeptides* | |

| | | | | |
|---|---|---|---|---|
| | | 30%-40%) | | |
| Human_7 | Human | Q Exactive (Fragmentation: SCE-HCD 20%-30%-40%) | PXD005931[7] | 6589 unique glycopeptides* |
| Human_8 | Human | Orbitrap Fusion Lumos (Fragmentation: SCE-HCD 31.5%-35%-38.5%) | PXD025455[8] | 606 unique glycopeptides* |
| Human_9 | Human | LTQ Orbitrap Elite (Fragmentation: SCE-HCD 20%-35%-50%) | PXD009716[9] | 40 unique glycopeptides* |
| Syn_1 | Synthesis glycopeptides | Orbitrap Fusion Tribrid (Fragmentation: SCE-HCD 20%-30%-40%) | PXD023980[5] | Sequence:<br>DLTHLJR<br>EEQFJSTFR<br>EEQYJSTYR<br>GHTLTLJFTR<br>LLNINPJK<br>RFJGSVSFFR<br>VQPFJVTQGK<br><br>Glycan:<br>Target:<br>(N(N(H(H(N(H)))(H(N(H(A)))))))<br>(N(N(H(H(N(H(A))))(H(N(H(A)))))))<br>Decoy:<br>(N(F)(F)(N(H(H(N(H)))(H(N(H))))))<br>(N(F)(N(H(H(N(H)))(H(N(F)(H))))))<br>(N(N(H(H(N(F)(H)))(H(N(F)(H))))))<br>(N(F)(F)(N(H(H(N(H)))(H(N(H(A)))))))<br>(N(F)(N(H(H(N(H)))(H(N(F)(H(A)))))))<br>(N(N(H(H(N(F)(H)))(H(N(F)(H(A)))))))<br>(N(F)(F)(N(H(H(N(F)(H)))(H(N(F)(H)))))) |
| Yeast_1 | Fission Yeast | Orbitrap Fusion Tribrid (Fragmentation: SCE-HCD 20%-30%-40%) | PXD023980[5] | 13,372 unique glycopeptides* |
| Yeast_2 | Fission Yeast | Orbitrap Fusion Tribrid (Fragmentation: SCE-HCD 20%-30%-40%) | PXD005565[1] | 16,809 unique glycopeptides* |

The term “unique glycopeptides” refers to the glycopeptides that exhibit distinct embedding patterns (glycan, peptide sequence, PTMs type, PTMs position, and precursor charge). The number of unique glycopeptides is based on the database searching software solution and filtration conditions detailed in **Methods**.

**Supplementary Table 2.** Nomenclature of monosaccharides used in this study.

| Name | Abbreviation | Character | Symbol |
|---|---|---|---|
| hexose | Hex | H | green circle |
| N-acetylhexosamine | HexNAc | N | blue square |
| N-acetylneuraminic acid | NeuAc | A | purple diamond |
| fucose | Fuc | F | red triangle |
| N-glycolylneuraminic acid | NeuGc | G | light blue diamond |

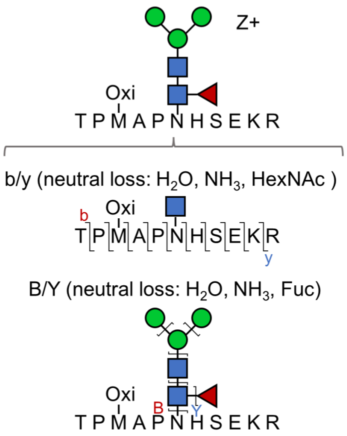


**Supplementary Figure 1.** The fragment types including neutral loss predicted by DeepGP.

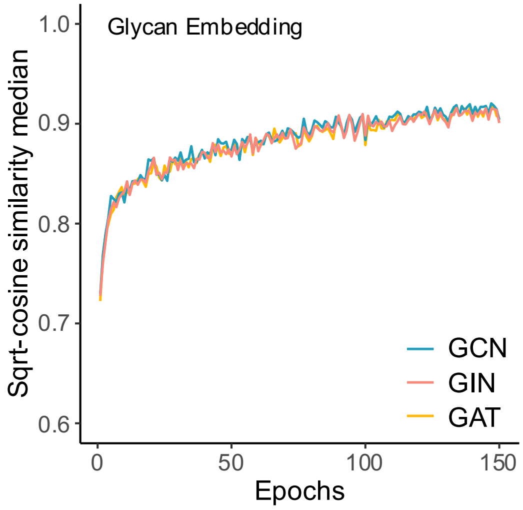


**Supplementary Figure 2. Comparison of GCN, GIN and GAT for glycan embedding.** Performance of GCN, GIN or GAT for glycan embedding during model training. The GNN used for B/Y ions intensity prediction was set as GIN. Model training was conducted with a batch size of 256, and a learning rate of $1\times10^{-4}$.

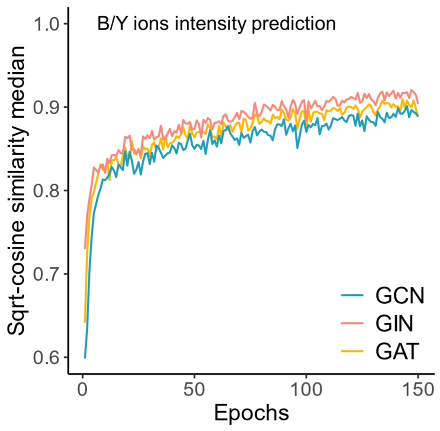


**Supplementary Figure 3. Comparison of GCN, GIN and GAT for B/Y ions intensity prediction.** Performance of GCN, GIN or GAT for B/Y ions intensity prediction during model training. The GNN used for glycan embedding was set as GCN. Model training was conducted with a batch size of 256, and a learning rate of $1\times10^{-4}$.

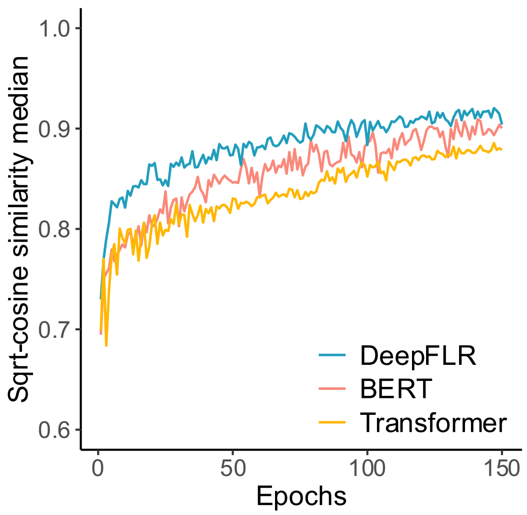


**Supplementary Figure 4. Comparison of DeepFLR, BERT and Transformer.** Performance of DeepFLR, BERT and Transformer during model training. Transformer means a Transformer model without pre-training, starting with randomly initialized parameters. Model training was conducted on with a batch size of 256, and a learning rate of $1\times10^{-4}$.

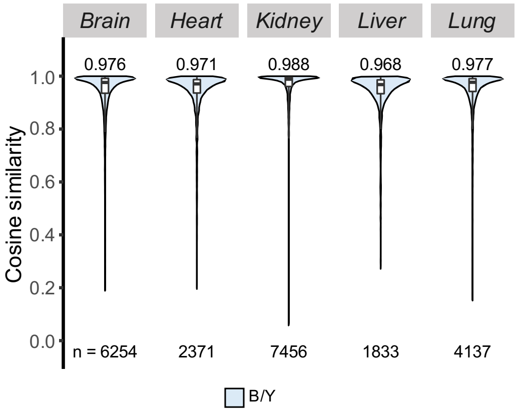


**Supplementary Figure 5. Performance of DeepGP in MS/MS spectra prediction for B/Y ions only.** The distribution of cosine similarity was computed between the predicted and experimental spectra for B/Y ions only. The medians are indicated. The boxes and whiskers show the quantiles and 95% percentiles, respectively. The numbers of spectra for test are indicated below each graph.

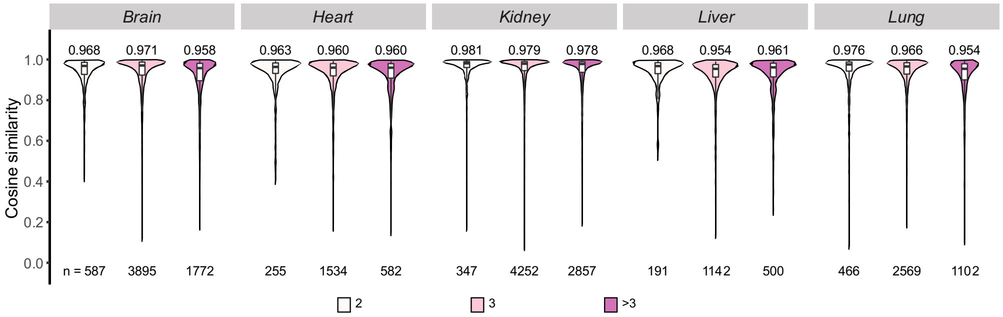


**Supplementary Figure 6. Performance of DeepGP in MS/MS spectra prediction across different charge states.** The distribution of cosine similarity was computed between the predicted and experimental spectra. 2: precursors with 2 charges; 3: precursors with 3 charges; >3: precursors with more than 3 charges. The medians are indicated. The boxes and whiskers show the quantiles and 95% percentiles, respectively. The numbers of spectra for test are indicated below each graph.

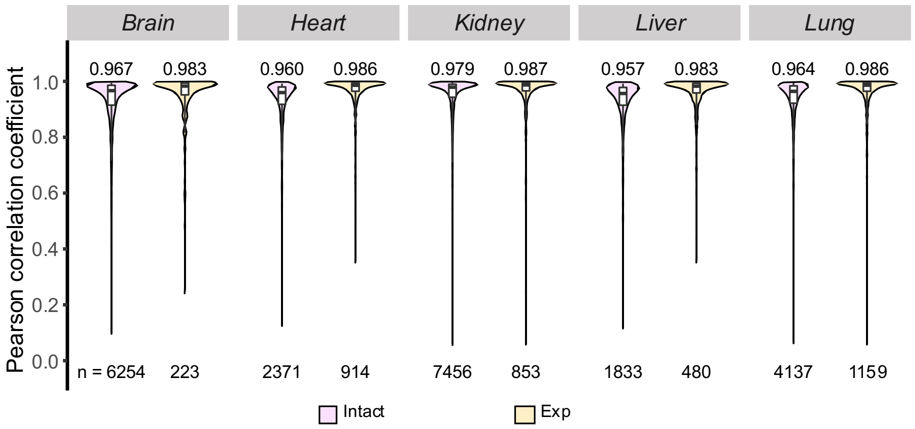


**Supplementary Figure 7. The distribution of Pearson correlation coefficient computed between the predicted and experimental spectra from the five mouse datasets.** Intact: Pearson correlation coefficient computed for all glycopeptides within the test datasets. Exp: Pearson correlation coefficient of repeatedly collected mass spectra in the training datasets and the test datasets of the same glycopeptides. The medians are indicated. The boxes and whiskers show the quantiles and 95% percentiles, respectively. The numbers of spectra used are indicated below each graph.

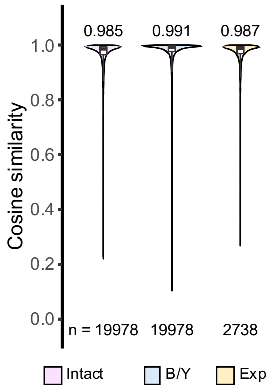


**Supplementary Figure 8. Performance of DeepGP in MS/MS spectra prediction trained with all the five datasets.** The distribution of cosine similarity computed between the predicted and experimental spectra for all the five mouse tissues data. Intact: cosine similarity computed for whole MS/MS spectra containing b/y and B/Y ions. B/Y: cosine similarity computed for B/Y ions only. Exp: cosine similarity of repeatedly collected mass spectra in the five mouse datasets of the same glycopeptides. The medians are indicated. The boxes and whiskers show the quantiles and 95% percentiles, respectively. The numbers of spectra for test are indicated below each graph.

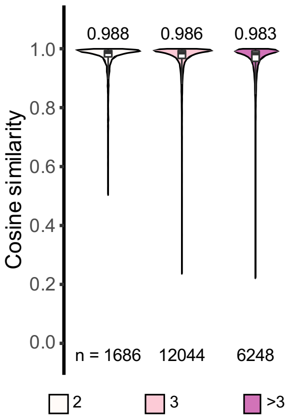


**Supplementary Figure 9. Performance of DeepGP in MS/MS spectra prediction trained with all the five datasets across different charge states.** The distribution of cosine similarity computed between the predicted and experimental spectra for all the five mouse tissues data. 2: precursors with 2 charges; 3: precursors with 3 charges; >3: precursors with more than 3 charges. The medians are indicated. The boxes and whiskers show the quantiles and 95% percentiles, respectively. The numbers of spectra for test are indicated below each graph.

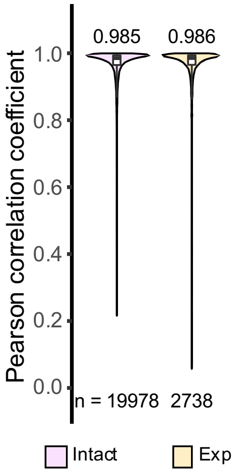


**Supplementary Figure 10. Performance of DeepGP in MS/MS spectra prediction trained with all the five datasets.** The distribution of Pearson correlation coefficient computed between the predicted and experimental spectra for all the five mouse tissues data. Intact: Pearson correlation coefficient computed for whole MS/MS spectra containing b/y and B/Y ions. Exp: Pearson correlation coefficient of repeatedly collected mass spectra in the five mouse datasets of the same glycopeptides. The medians are indicated. The boxes and whiskers show the quantiles and 95% percentiles, respectively. The numbers of spectra used are indicated below each graph.

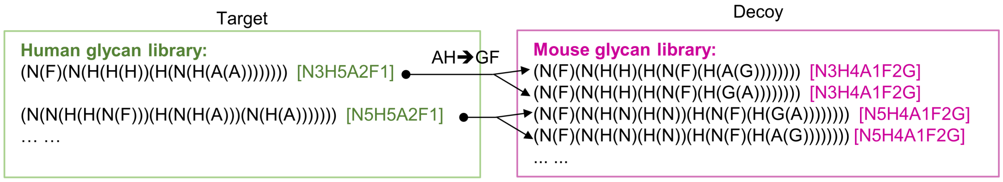


**Supplementary Figure 11.** Generation of decoy glycans for the two human datasets. The decoy glycan for human sample was generated by substituting NeuAc and Hex with NeuGc and Fuc, using the glycan available in the mouse glycan library.

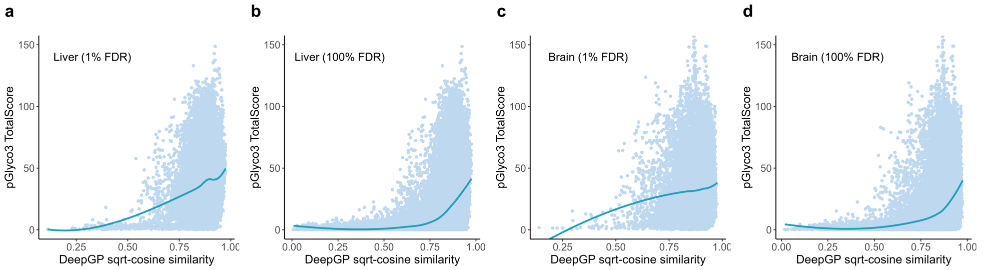


**Supplementary Figure 12. The correlation between DeepGP sqrt-cosine similarity and pGlyco3 TotalScore for (a) mouse liver data at 1% FDR (b) mouse liver data at 100% FDR (c) mouse brain data at 1% FDR and (d) mouse brain data at 100% FDR.** Each point represents a data observation. The locally estimated scatterplot smoothing (LOESS) line is also plotted to highlight the local regression fit.

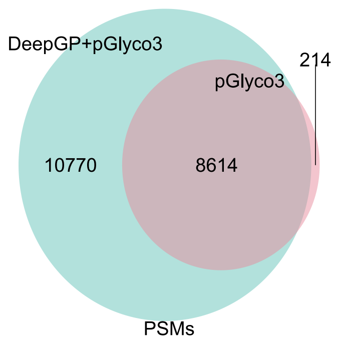


**Supplementary Figure 13.** Venn diagram of the number of PSMs identified by DeepGP+pGlyco3 and pGlyco3 alone for Yeast_1 at the decoy ratio of 5%.

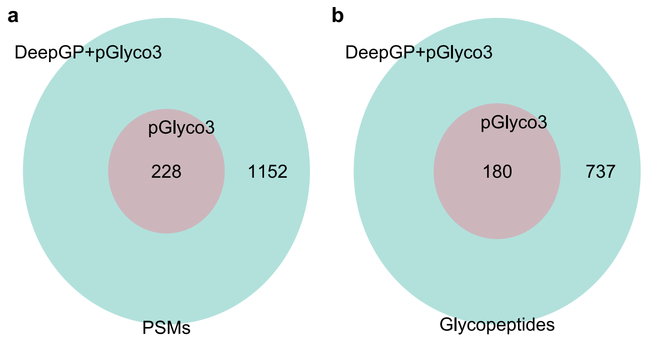


**Supplementary Figure 14.** Venn diagram of the number of **(a)** PSMs and **(b)** glycopeptides identified by DeepGP+pGlyco3 and pGlyco3 alone for Yeast_1 at the decoy ratio of 1%.

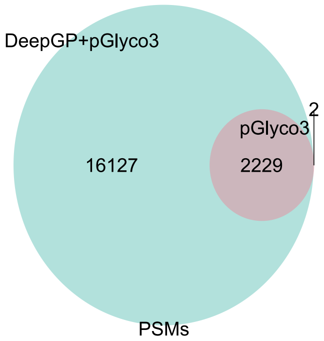


**Supplementary Figure 15.** Venn diagram of the number of PSMs identified by DeepGP+pGlyco3 and pGlyco3 alone for Yeast_2 at the decoy ratio of 5%.

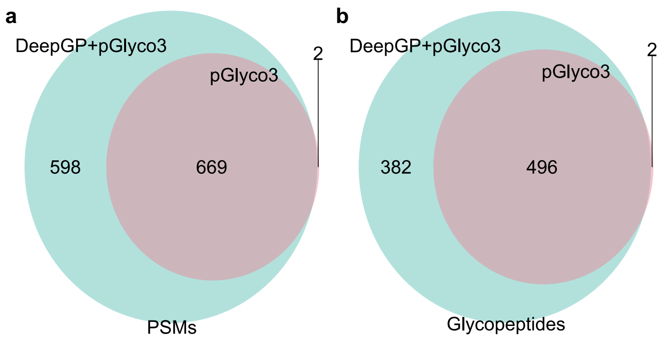


**Supplementary Figure 16.** Venn diagram of the number of **(a)** PSMs and **(b)** glycopeptides identified by DeepGP+pGlyco3 and pGlyco3 alone for Yeast_2 at the decoy ratio of 1%.

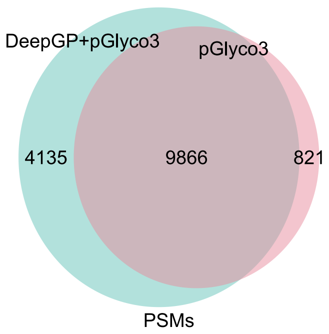


**Supplementary Fig. 17.** Venn diagram of the number of PSMs identified by DeepGP+pGlyco3 and pGlyco3 alone for Mouse_Brain at the decoy ratio of 5%.

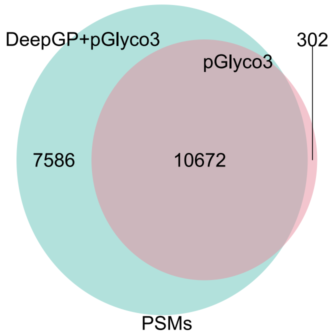


**Supplementary Fig. 18.** Venn diagram of the number of PSMs identified by DeepGP+pGlyco3 and pGlyco3 alone for Mouse_Liver at the decoy ratio of 5%.
